# Ultra-low Input Circulating Tumor DNA Detection by MED-Amp in Early-Stage Pancreatic Cancer

**DOI:** 10.1101/2021.03.28.437388

**Authors:** Erica D. Pratt, David B. Zhen, Robert W. Cowan, Heather Cameron, Kara Schradle, Sara L. Manning, Valerie Gunchick, Diane M. Simeone, Vaibhav Sahai, Andrew D. Rhim

**Author notes:** **Corresponding author:** Erica D. Pratt, University of Minnesota Twin Cities, 420 Washington Ave SE, MCB 7-194, Minnesota, MN 55455.

## Abstract

**Purpose:** The clinical utility of circulating tumor DNA (ctDNA) has been shown in advanced pancreatic ductal adenocarcinoma (PDA). However, diagnostic sensitivity of many ctDNA assays is low in resectable and locally advanced disease, where tumor burden is substantially lower. We have previously described Multiplex Enrichment using Droplet Pre-Amplification (MED-Amp), a multiplexed panel for the detection of the most common oncogenic *KRAS* mutations in PDA. In this study, we aimed to assess the diagnostic sensitivity of MED-Amp for detection of rare mutant alleles present in the plasma of patients with localized PDA.

**Experimental Design:** We retrospectively analyzed ninety-eight plasma samples from 51 patients with various stages of localized disease. For comparison, we measured ctDNA levels in 20 additional patients with metastatic PDA. The MED-Amp assay was used to measure the abundance of the four most common *KRAS* codon 12 mutations (G12C/D/R/V). We correlated the presence and quantity of ctDNA with overall survival (OS) as well as progression-free survival (PFS). Using serial plasma draws, we also assessed the relationship between changes in ctDNA allelic frequency and progression.

**Results:** *KRAS*-positive ctDNA was detected in 52.9% of localized PDA and 75% of metastatic samples tested using DNA inputs as low as 2 ng. As previously reported, the presence of *KRAS* mutant ctDNA was correlated with worse OS for all disease stages (p = 0.02). In patients with localized PDA high ctDNA levels also correlated with significantly worse median OS (533 days vs 1090 days) and PFS (192 days vs 787 days). We also studied a small cohort of serial plasma draws to observe the relationship between ctDNA fold change and PFS. We found 83% of patients with increased fold change in mutant *KRAS* experienced disease progression (n=6). In contrast, 75% (n=4) of patients with decreased fold change remained disease-free (p=0.03).

**Conclusions:** MED-Amp is a flexible and cost-effective approach for measurement of ctDNA in patients with localized cancer. Though this study focused on *KRAS* mutation detection, this assay could be adapted for a number of common oncogenic alterations.

**Statement of translational relevance:** Only 25% of pancreatic ductal adenocarcinoma (PDA) patients with localized disease survive five years post-resection. It is hypothesized PDA undergoes dissemination at the earliest stages of tumor formation, driving formation of occult metastases which go undetected using conventional screening methods. Development of high specificity, high sensitivity biomarkers is critical to improving patient mortality. Circulating tumor DNA (ctDNA) has gained increasing acceptance as a non-invasive prognostic in metastatic disease. However, the sensitivity of most targeted ctDNA assays precludes reliable detection of localized and resected disease. Here, we present a digital droplet PCR assay for multiplexed enrichment and detection of *KRAS* mutations, the most commonly mutated oncogene in PDA. This assay preserves ctDNA allelic frequency in the original sample, while increasing the molecular signal over 50-fold. This study shows the ctDNA has potential diagnostic value in early-stage PDA, and that digital preenrichment of cell-free DNA increases overall assay sensitivity without sacrificing specificity.

## Introduction

Pancreatic adenocarcinoma (PDA) is projected to be the second leading cause of cancer deaths by 2030 and has a five-year survival rate of only 9% (1,2). While high patient mortality is driven primarily by late stage diagnosis, patients with localized disease who undergo curative resections still have a five-year survival rate of only 25% (3). Based on these data, it has been suggested that tumor dissemination begins at the earliest stages of PDA development, and research from genetically engineered mouse models of PDA support this hypothesis (4). These occult metastases could go undetected using conventional screening methods, thereby fueling cancer progression in patients clinically staged as having localized disease. A surrogate marker of cancer progression could provide invaluable information during therapeutic decision-making. However, there are no clinically validated biomarkers with the required specificity or sensitivity for surveillance of localized PDA.

Research in metastatic disease has shown the presence of circulating tumor DNA (ctDNA) is correlated with tumor load and patient survival, as well as chemotherapeutic efficacy in late-stage PDA (5,6). However, detection remains challenging in localized disease, where ctDNA is often present at much lower levels. Bettegowda and colleagues quantified the relationship between disease stage and ctDNA abundance in PDA and 15 other cancers using a combination of next-generation sequencing (NGS) with unique molecular identifiers (SafeSeqS), and digital PCR followed by flow cytometry (BEAMing) (7). Median ctDNA burden in localized PDA was 1.2 mutant alleles per milliliter of plasma, nearly 10 times lower than metastatic samples. Recent studies have used patient medical history in combination with NGS for accurate detection of PDA, as well as a variety of epithelial cancers, achieving a diagnostic sensitivity of 72% (8). However, NGS typically requires high concentrations of cell-free DNA (cfDNA) for library construction and can have multi-day turnaround times due to sequencing, alignment, variant filtering, and data interpretation. Additionally, the amplification and sequencing error rates inherent to NGS platforms decrease call accuracy when variant allelic frequencies (VAFs) fall below 0.1%.

Digital PCR (dPCR) is a cost-effective alternative, relying on physical partitioning samples into individual microfluidic reaction volumes to increase PCR efficiency. DNA template is then simultaneously amplified and then detected using hydrolysis-based fluorescent probes in a single PCR reaction. The total number of partitions, rather than quantity of mutant alleles, determines the theoretical lower limit of detection (LLOD). This dramatically increases diagnostic sensitivity for molecular targets as rare as 1 mutant allele in 1,000,000 wild-type copies (0.001%) (9). dPCR also allows for absolute quantification of mutant DNA, enabling investigation of ctDNA and its relationship with cancer progression (8,10–16). Prior research in advanced and localized cancer has shown ctDNA VAF measured by dPCR is also highly concordant with NGS (12,17,18). These qualities make dPCR ideal for assaying small panels of known mutations. However, dPCR has yet to realize its full potential for the detection of localized disease, where both cfDNA concentrations and mutant allele frequencies are low. Zhang and colleagues reported dPCR assay sensitivity decreased 4-fold when cfDNA total input was under 5 ng (19). Brychta et al. reported that *KRAS* ctDNA was 50% less likely to be detected if median cfDNA input was under 20 ng (13). Yet multiple studies have reported up to 25%-30% of early-stage plasma samples have cfDNA concentrations under 10 ng/mL (20,21). Large volume plasma draws to recover more cfDNA can increase analytical sensitivity; however, this is often impractical.

We have previously reported on Multiplex Enrichment using Droplet Pre-Amplification (MED-Amp) as an effective method to increase assay sensitivity in low cfDNA input, low ctDNA abundance liquid biopsy samples (22). Using our strategy, DNA template undergoes single molecule emulsification followed by a short PCR preamplification step to uniformly increase total signal over 50-fold. The amplified template is purified and then quantified by conventional dPCR. Single molecule emulsification circumvents known PCR bias against rare alleles, and hard-to-amplify sequences, such as mutations in *TP53* and *KRAS* (23). MED-Amp also does not rely on selective digestion of wild-type (WT) template to enhance mutant (MT) signal, retaining valuable mutation VAF information. Here, we use our MED-Amp assay for ctDNA detection and quantification in a retrospective analysis of 91 plasma samples from seventy-one PDA patients with localized and advanced PDA. We assessed assay sensitivity for cfDNA inputs as low as 2 ng, which can preclude successful NGS library preparation or dPCR detection. We also aimed to assess the prognostic value of ctDNA for patients with early-stage PDA.

## Materials and Methods

### Patient Characteristics

All samples were collected with Institutional Review Board approval (HUM25339) at University of Michigan and under compliance with HIPPAA guidelines. We analyzed ninety-eight plasma samples from seventy-one pancreatic cancer patients, and twenty-five additional samples from non-cancer controls undergoing routine colonoscopy. Samples were collected from patients with resectable (n=17), borderline (n=16), locally advanced (n=18), and metastatic (n=20) disease.

### Patient Plasma Isolation and DNA Extraction

Patient blood samples were drawn in either Streck or EDTA tubes and were processed within 30 minutes of collection. Samples were spun for 10 minutes at 820 xg at 4°C. The plasma supernatant was extracted via pipette and underwent a second spin at 16,000 xg for 10 minutes at 4°C. The supernatant was aliquoted in 1 mL volumes in 1.5 mL Eppendorf tubes and stored at −80°C until further processing. Matching buffy coat was also collected and stored at −80°C. cfDNA was isolated from 1.5-2 mL of plasma using the QIAmp Circulating Nucleic Acid Kit as specified by the manufacturer. DNA was eluted in 150 μL of AVE buffer (RNase-free water with 0.04% sodium azide), and then concentrated using ethanol precipitation as described previously (22) in a final volume of 15 μL. Median total cfDNA recovered was 9.2 ng (IQR: 6.3-14.7 ng). The mean time from plasma storage to DNA isolation and processing was 28 months.

### Preamplification Using Droplet-based PCR

Fourteen microliters of precipitated cfDNA sample was used to generate 50 μL volume reaction mixes for emulsification. The RainDance Source digital PCR system was used to partition the reaction mix into 5 pL volume droplets prior to preamplification using Q5 Hot-Start High Fidelity polymerase as described previously (10). Amplified template was de-emulsified and underwent PCR cleanup prior to re-partitioning with *KRAS* WT and MT-specific TaqMan probes. Droplets were processed using the RainDance Sense digital PCR system, using a VIC reporter to label *KRAS* WT and FAM for *KRAS* G12C, G12D, G12R, and G12V mutations. Resulting populations were gated with a standardized template using RainDance Analyst II™ software. Variant allelic frequency (VAF) was calculated by the number of FAM positive droplets divided by the sum of FAM and VIC droplets. Samples where the MT droplet count was below the assay LLOD (33 FAM-positive droplets) were classified negative for *KRAS* mutations and assigned a VAF of 0%.

### Statistical Analysis

All data analyses were performed using R 3.4.4, a multi-platform open-source language and software for statistical computing (24). Continuous variables were reported as median and interquartile range (IQR) and compared using the Mann-Whitney-Wilcoxon test or Kruskal-Wallis test unless otherwise noted. Results significant by Kruskal-Wallis were subjected to a Dunn’s test post-hoc with Benjamin-Hochberg correction for multiple comparisons using the ‘PMCRplus’ R package. Categorical count data was compared using either the Chi-Squared Test without continuity correction or Fisher’s Exact Test. The Tarone-Ware variant of the log-rank test was used to quantify differences in trends between Kaplan-Meier survival curves. Survival analyses were performed using a combination of R packages ‘survminer’ and ‘survival’.

## Results

### Patient Characteristics

Ninety-eight plasma samples from seventy-one pancreatic cancer patients were selected for retrospective analysis. The localized PDA cohort was comprised of resectable (n=17), borderline (n=16), and locally advanced (n=18) patients. We compared results from localized PDA samples with those from 20 patients with metastatic PDA. The patients were 47% male, 53% female, with a median age of 65 yrs. Tumors were most frequently found at the head of the pancreas (52%), followed by the body (14%), tail (10%), neck (5%), or a combination (19%). There were no statistically significant differences in patient cohorts between the localized and metastatic patient groups. Patient details and characteristics are outlined in Table 1, and indepth clinical information is available in Table S1. Our PDA cohort included patients who had undergone neoadjuvant (32%) or adjuvant chemotherapy (34%), either alone, or in combination with radiation (34%). At the time of analysis, 53% (n=27) of PDA patients had died. For seven patients, prior medical records were unavailable due to treatment at an external institution. Of the localized PDA patients where complete medical history was available (n=34), 50% progressed in a median time of 263 days. Control samples were drawn from twenty-five patients undergoing routine colonoscopy. The control group was younger than the PDA cohort, with a median age of 54 yrs (p < 0.001). Nineteen control patients had a prior history of benign polyps or inflammation, and six had no prior history of cancer, colorectal polyps, or inflammatory conditions (Table S2).

**Table 1:** PDA Patient Characteristics.

|  | Total | Resectable | Borderline | Locally<br>Advanced | Metastatic |
| --- | --- | --- | --- | --- | --- |
| Characteristics | n=71 | n=17 | n=16 | n=18 | n=20 |
| Age, median (IQR), yrs | 65 (60-73) | 70 (63-75) | 65 (57-68) | 64 (61-74) | 65 (59-68) |
| Gender, no. (%) |  |  |  |  |  |
| Female | 39 (53.4) | 7 (43.8) | 10 (55.56) | 13 (68.42) | 9 (45) |
| Male | 34 (46.6) | 9 (56.3) | 8 (44.44) | 6 (31.58) | 11 (55) |
| Tumor Location, no. (%) |  |  |  |  |  |
| Head | 38 (52.1) | 12 (75) | 8 (44.4) | 9 (47.4) | 9 (45) |
| Neck | 4 (5.5) | 0 (0) | 2 (11.1) | 2 (10.5) | 5 (25) |
| Body | 10 (13.7) | 1 (6.3) | 2 (11.1) | 4 (21.1) | 3 (15) |
| Tail | 7 (9.6) | 3 (18.8) | 1 (5.6) | 0 (0) | 3 (15) |
| Combination | 14 (19.2) | 0 (0) | 5 (27.8) | 4 (21.1) | 0 (0) |

**PDA:** Pancreatic Adenocarcinoma

Blood was collected from each patient, and plasma and buffy coat fractions were extracted and stored at −80°C until further use. A maximum of 2 mL of plasma from each patient underwent MED-Amp as previously described (Fig. 1) (22). Measured VAF was correlated with tumor size (Pearson r = 0.3, p = 0.05, Figure S1A), consistent with prior observations (25). There was no dependence between DNA input and measured VAF, or sample age and total

**Figure 1.**
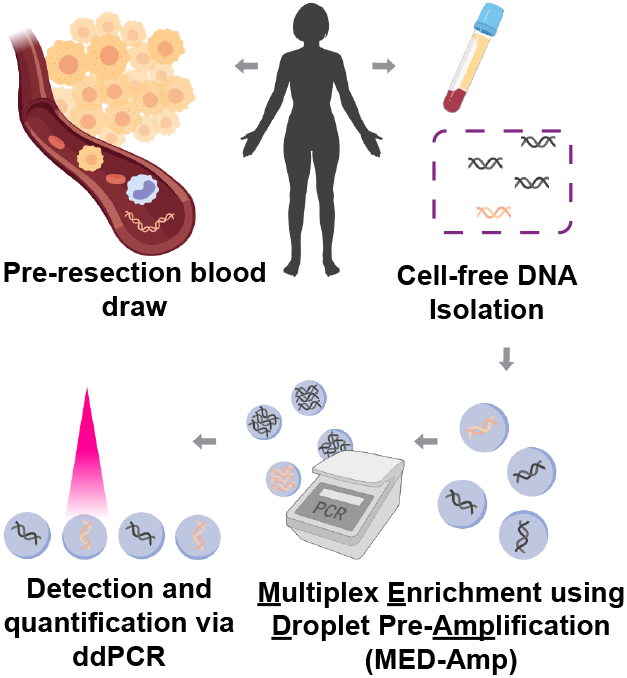
Overview of ctDNA analysis. Cell-free DNA was isolated from ninty-eight plasma samples from seventy-one pancreatic cancer patients. Targeted digital preamplification of a 90 bp region in *KRAS* codon 12 was performed as described previously (22). The enriched amplicon was then analyzed for *KRAS* mutations using dPCR.

DNA recovered (Figure S1B & C). Average cfDNA levels were significantly higher in cancer patients (14.4 ng) versus non-PDA controls (8.4 ng, p = 0.03). However, there was no statistically significant difference in cfDNA concentration between patients with localized versus metastatic disease (p = 0.79, Figure 2A).

**Figure 2.**
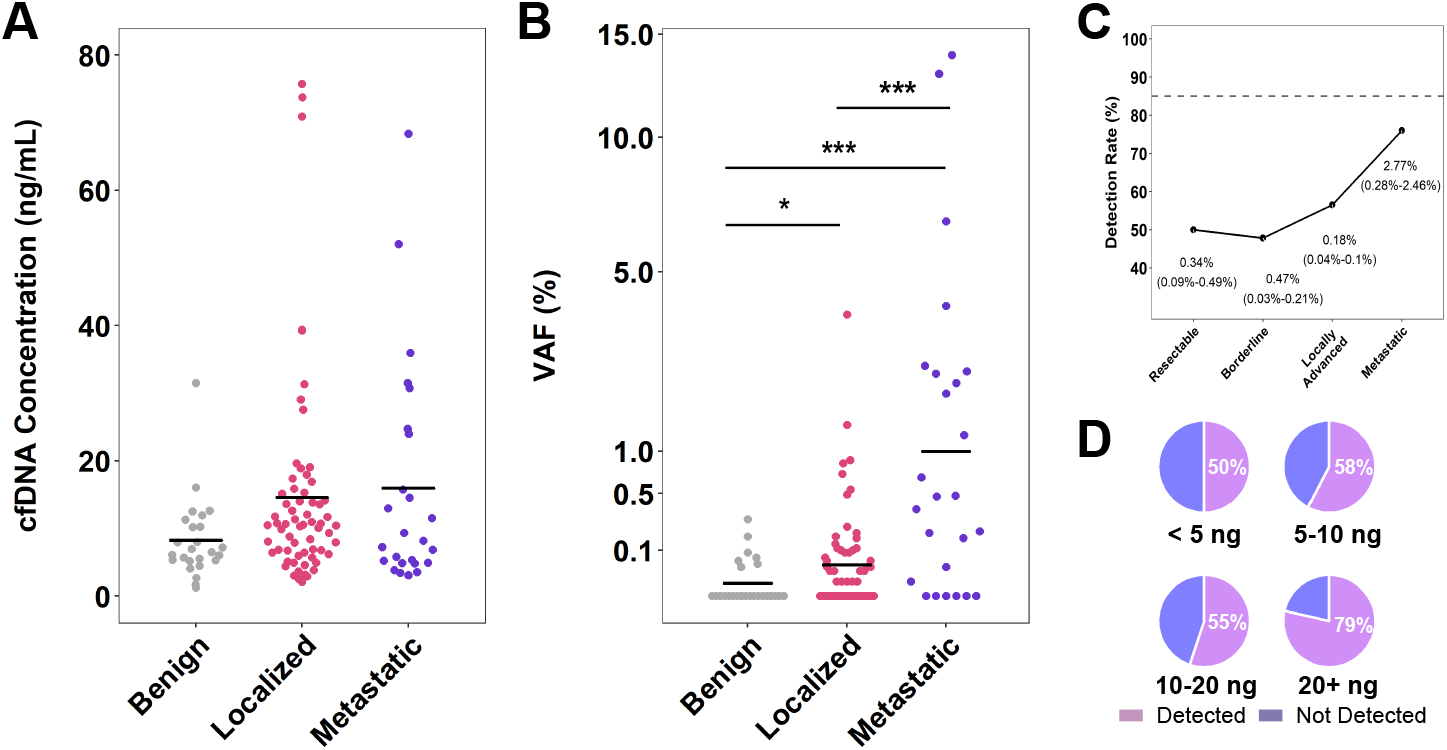
MED-Amp detection of *KRAS* mutant ctDNA in early-stage and metastatic. A) cfDNA concentration (ng/mL) increases with cancer stage. B) Mutant *KRAS* variant allelic frequency (VAF) in metastatic and localized PDA co horts compared to controls. * p < 0.05 *** p < 0.001. C) Sensitivity of *KRAS* ctDNA detection increases with overall tumor burden. Median VAF and IQR reported for each disease stage. D) MED-Amp analytical sensitivity for *KRAS* as a function of total cfDNA input.

### *KRAS* ctDNA levels in patients with localized and metastatic PDA

Using MED-Amp, we measured *KRAS* ctDNA levels in each patient group. *KRAS* abundance was significantly higher than in both localized (p=0.044) and metastatic samples (p < 0.001 compared to control samples (Figure 2B). As expected, average metastatic patient ctDNA VAFs was 9-fold higher than localized cases (p < 0.001). *KRAS* ctDNA was successfully detected in 52.9% of patients with localized PDA, with minimal variation across cohorts: resectable (n=9/17, 52.9%), borderline (n=7/16, 43.8%), locally advanced (n=11/18, 61.1%). Similar to previous reports, 75% (n=15/20) of metastatic PDA patients tested positive for *KRAS* mutant ctDNA. Assay sensitivity was higher in the metastatic cohort, but this difference was not statistically significant (p = 0.109). Overall, we successfully detected *KRAS* ctDNA in at least one plasma sample for 42 of 71 (59.2%) PDA patients tested (Figure 2C).

Our control cohort was predominantly patients with prior medical history of chronic inflammation or colorectal adenomas (76%). Colorectal adenomas are common in aging populations (≥ 25%) and may be a confounding factor during molecular testing (26). Prior studies have reported *KRAS* codon 12 mutation-containing circulating DNA can be found in up to a third of patients with polyps but no evidence of malignancy (27). We first analyzed the rate of mutant *KRAS* detection in our patients with no history of adenomas or inflammation. Assay specificity was quite high, with 83% (n=5/6) of samples testing negative for *KRAS* mutations. As expected, mutant *KRAS* levels were higher in patients with prior history of adenomas or inflammation, with 6 out of 19 (32%) testing positive for *KRAS* mutations.

### MED-Amp performance at low cfDNA inputs

We next evaluated MED-Amp sensitivity as a function of cfDNA input. In our data set, over half of the cfDNA inputs were below 10 ng, and 20% were less than 5 ng (Figure S2A). The median cfDNA input for our assay was 8.5 ng, or approximately 2500 genomic equivalents. MED-Amp sensitivity was robust across all ranges tested, increasing slightly at DNA inputs recommended by RainDance Technologies for optimal performance (Figure 2D). We successfully detected *KRAS* ctDNA mutations at VAF of 0.04 % in cfDNA concentrations as low as 5 ng (~1 mutant copy). We also assessed assay repeatability by performing two independent MED-Amp experiments on six representative DNA samples from patients with benign, resectable, and locally advanced disease. There was 100% concordance between samples on presence/absence of ctDNA. Measured VAFs were highly correlated with an R^2^ value of 0.94 (Figure S2B).

### Prognostic value of ctDNA for patient survival

We first assessed *KRAS* ctDNA correlation with overall survival (OS) or progression-free survival (PFS). Presence of *KRAS* mutations was predictive with patient death regardless of disease stage (p = 0.02) but did not predict disease progression (p = 0.38). Based on these results, we stratified ctDNA VAF into “high” and “low” groups based on OS for the localized and metastatic cohorts using maximally selected ranked statistics. Low ctDNA VAF was predictive of better survival outcomes in both localized and metastatic patient cohorts in univariate analyses (HR: 3.2, 95% CI: 1.4-7.3, p=0.009) (Table 2). The other predictive variable was prior radiation therapy, possibly driven by the inclusion of borderline PDA patients undergoing radiotherapy to shrink tumor volume. ctDNA remained an independent predictor of patient OS after multivariate analysis (HR: 3.88, 95% CI: 1.34-11.22, p = 0.012). Low ctDNA levels correlated with a near doubling of median survival time in localized (533 vs 1090 days, p=0.0089) and metastatic disease (334 vs 735 days, p = 0.0342) (Figure 3A & B).

**Figure 3.**
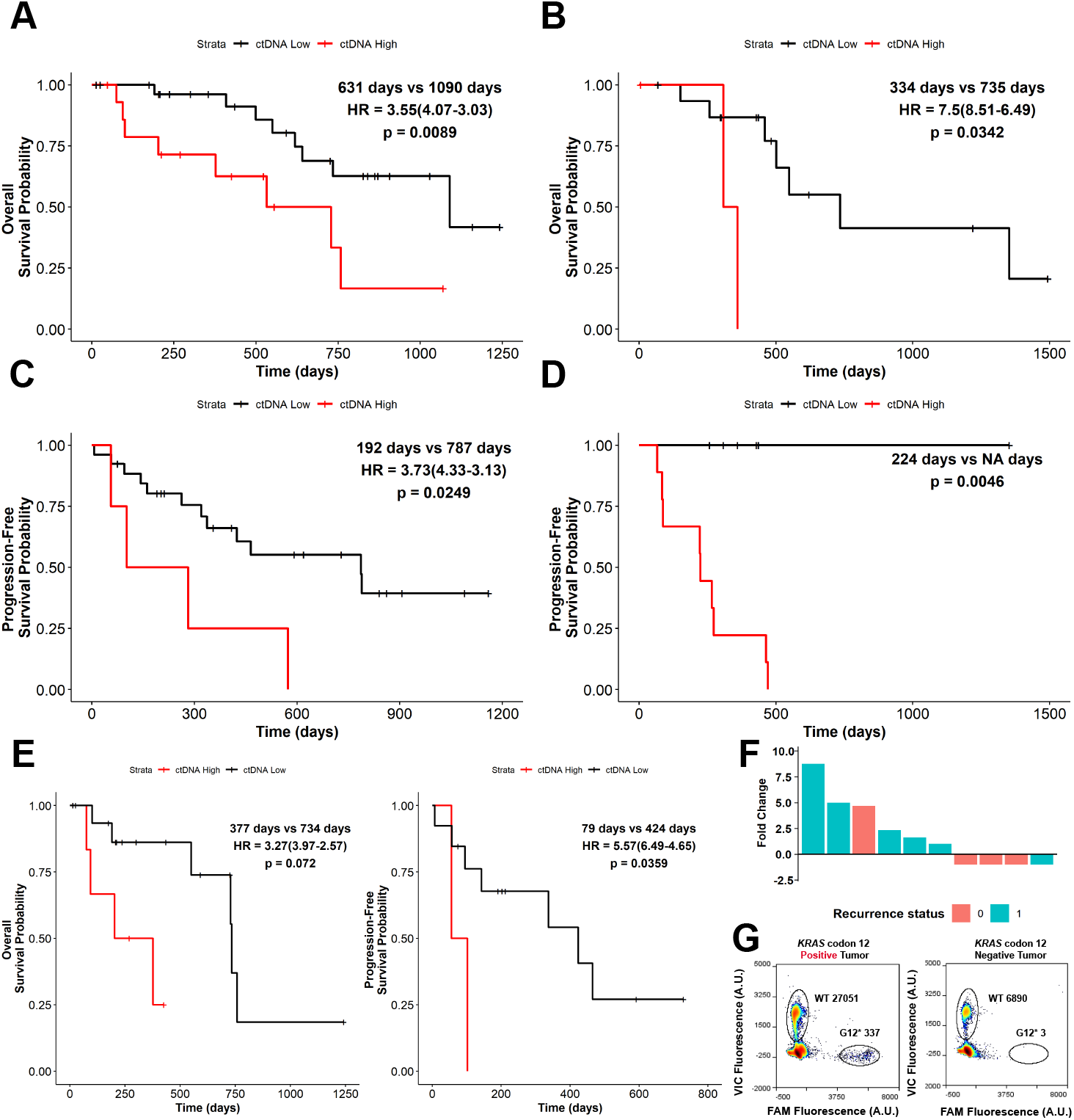
Kaplan-Meier analysis of overall survival (OS) and progression-free survival (PFS). Stratification of patients by ctDNA prevalence reveals patients with higher ctDNA burden have significantly lower OS in localized (A) and metastatic (B) PDA. High ctDNA frequency is also predictive of reduced time to progression in localized (C) and metastatic (D) disease. Patients with lower mutant *KRAS* levels lived almost twice as long and had longer time to relapse (E). Analysis of VAF fold-change (FC) between serial blood draws (F) reveals fold change direction correlates with PDA progression (p=0.03, one-sided Chi-Squared Test). Comparison of MED-Amp with mutational analysis of tissue showed ctDNA mutations reflected those found in the primary tumor (G).

**Table 2:**
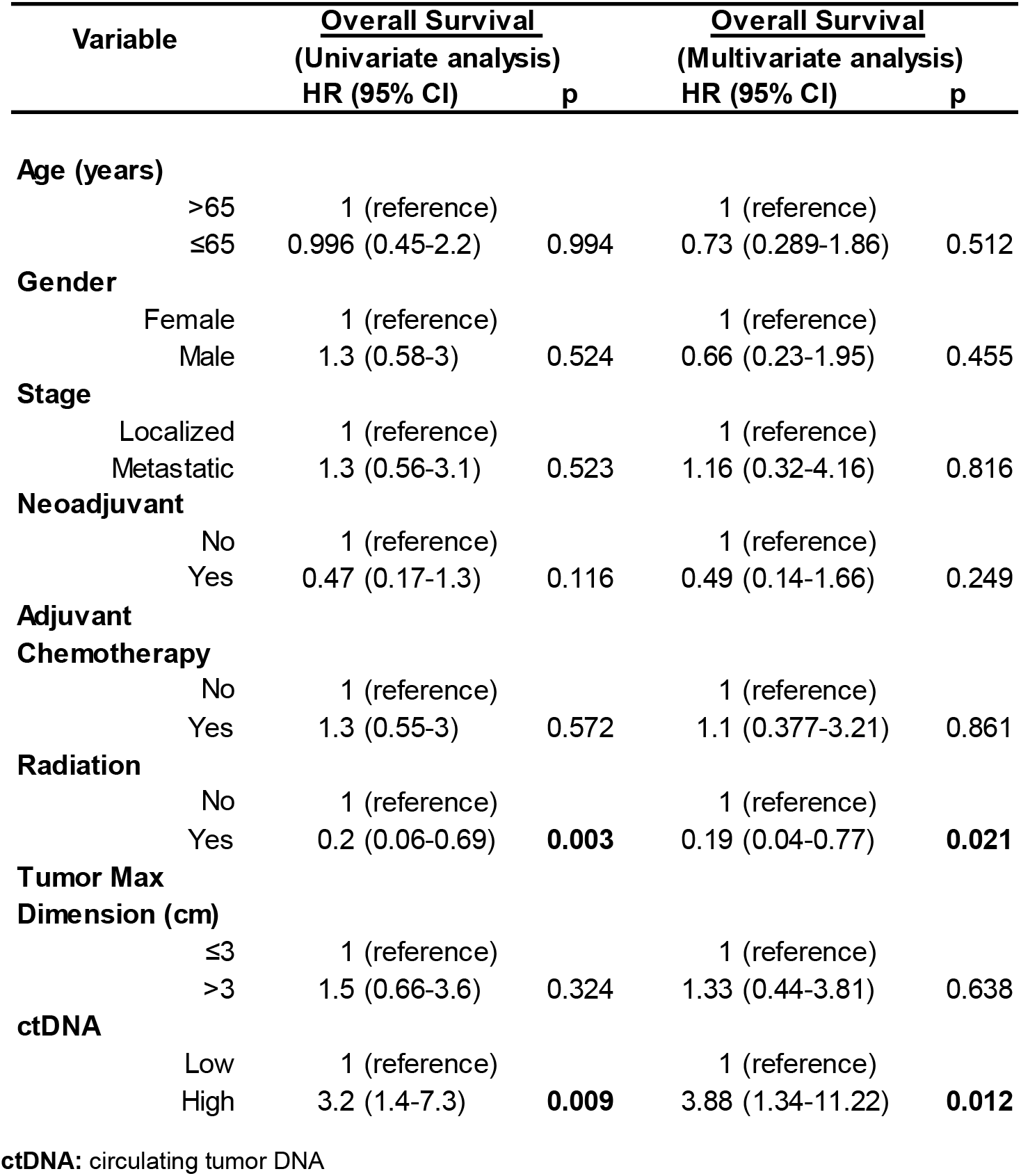
Uni- and Multivariate Analyses.

| Variable | <u>Overall Survival</u> |  | <u>Overall Survival</u> |  |
| --- | --- | --- | --- | --- |
|  | (Univariate analysis) |  | (Multivariate analysis) |  |
|  | HR (95% CI) | p | HR (95% CI) | p |
| **Age (years)** |  |  |  |  |
| >65 | 1 (reference) |  | 1 (reference) |  |
| ≤65 | 0.996 (0.45-2.2) | 0.994 | 0.73 (0.289-1.86) | 0.512 |
| **Gender** |  |  |  |  |
| Female | 1 (reference) |  | 1 (reference) |  |
| Male | 1.3 (0.58-3) | 0.524 | 0.66 (0.23-1.95) | 0.455 |
| **Stage** |  |  |  |  |
| Localized | 1 (reference) |  | 1 (reference) |  |
| Metastatic | 1.3 (0.56-3.1) | 0.523 | 1.16 (0.32-4.16) | 0.816 |
| **Neoadjuvant** |  |  |  |  |
| No | 1 (reference) |  | 1 (reference) |  |
| Yes | 0.47 (0.17-1.3) | 0.116 | 0.49 (0.14-1.66) | 0.249 |
| **Adjuvant** |  |  |  |  |
| **Chemotherapy** |  |  |  |  |
| No | 1 (reference) |  | 1 (reference) |  |
| Yes | 1.3 (0.55-3) | 0.572 | 1.1 (0.377-3.21) | 0.861 |
| **Radiation** |  |  |  |  |
| No | 1 (reference) |  | 1 (reference) |  |
| Yes | 0.2 (0.06-0.69) | **0.003** | 0.19 (0.04-0.77) | **0.021** |
| **Tumor Max** |  |  |  |  |
| **Dimension (cm)** |  |  |  |  |
| ≤3 | 1 (reference) |  | 1 (reference) |  |
| >3 | 1.5 (0.66-3.6) | 0.324 | 1.33 (0.44-3.81) | 0.638 |
| **ctDNA** |  |  |  |  |
| Low | 1 (reference) |  | 1 (reference) |  |
| High | 3.2 (1.4-7.3) | **0.009** | 3.88 (1.34-11.22) | **0.012** |

**ctDNA:** circulating tumor DNA

### Prognostic value of ctDNA for disease progression

ctDNA stratification for patient PFS was performed as described above, and high ctDNA was also found to also be predictive of disease progression (Figure 3C & D). We further investigated the prognostic ability of ctDNA specifically in non-metastatic PDA. As with OS, low ctDNA VAF was correlated with delayed median time to progression (787 vs 192 days, p = 0.025). Abundance of ctDNA was also correlated with OS and PFS in patients with un-resected localized disease (Figure 3E). In the four patients who had resections and experienced no recurrence within the time period of interest, 100% had undetectable ctDNA prior to surgery. A quarter of resected patients who eventually experienced disease recurrence (n=8) had detectable ctDNA pre-surgery. However, ctDNA presence was not predictive in patients who went on to have a resection (Figure S3A & B).

### Serial measurements of ctDNA and disease progression

Serial blood draws were obtained from 11 PDA patients, seven of whom had localized cancer. Median time between blood draws was 112 days (IQR: 98-129 days). The median time to cancer progression was 337 days. We measured relative fold change (FC) in ctDNA levels, defined as (VAFdraw #2 ÷ VAFdraw #1) - 1. We considered an absolute FC change of greater than 0.5 as a true shift in ctDNA burden. 10 out of 11 patients experienced |FC| greater than 0.5, most of which experiencing a positive FC between draws (n=6). Increased allele FC was positively correlated with cancer progression (p = 0.03) (Figure 3F).

### Correlation between tumor and ctDNA genotype

Data from genomic profiling of the primary tumor was available for 4 of the 71 patients assayed. Patients 278 and 427 had confirmed *KRAS* p.G12D mutations, while patients 684 and 814 both had *KRAS* p.Q61H mutations. Patient 684 had an additional copy number increase of the *KRAS* gene. Patients 278, 427, and 684 had metastatic PDA while patient 814 had resectable disease. The median time from diagnosis to plasma draw was 189 days (IQR: 45-353 days). As expected, plasma from patients 278 and 427 tested positive for *KRAS* codon 12 mutations. Patients 684 and 814 did not have *KRAS* codon 12 mutations, and their plasma samples were also *KRAS* codon 12 negative (Figure 3G). These data suggest that ctDNA accurately reflected the original tumor genotype.

## Discussion

In our prior work, we showed that PCR preamplification of single-molecule emulsions prior to measurement by dPCR greatly enhances assay sensitivity without compromising the quantitative advantages of dPCR (22). Here, we extend this strategy to show assay performance is uncompromised by the low cfDNA inputs common with non-metastatic cancer samples. This is a key advancement, as DNA input is a crucial pre-analytical factor that critically affects sensitivity of PCR-reliant methodologies (28,29). We successfully detected *KRAS* mutant ctDNA in 52.9% of non-metastatic and 75% of metastatic patients. We observed no statistically significant drop in assay sensitivity between resectable and locally advanced PDA, as has been reported in other studies (8,10,13). Our results are consistent with other molecular assays in localized PDA, which have reported sensitivities ranging from 31% to 62% (7,11,12,14,30).

In agreement with prior studies, we observed that *KRAS* mutant ctDNA was prognostic for overall and progression-free survival in patients with non-metastatic PDA. Survival time of patients with early-stage PDA and elevated ctDNA levels was half that of patients with low ctDNA (533 vs 1090 days, p=0.0089), and progression-free survival was similarly reduced (192 vs 787 days, p=0.025). All patients with undetectable ctDNA prior to resection experienced no recurrence, while a quarter of patients with detectable ctDNA eventually progressed. Serial monitoring of ctDNA dynamics was also predictive of PDA progression. Eighty-three percent of patients who experienced an increase fold-change in ctDNA between blood draws experienced relapse, while 75% of patients with decrease fold-change did not (p=0.03). When molecular profiling information for the primary tumor was available, there was 100% concordance between tumor and ctDNA genotype. Our initial results support a growing body of evidence suggesting ctDNA is a suitable prognostic of patient survival in localized PDA.

Several limitations to our study should be acknowledged. First, our study is retrospective, with a small total number of patients (n=71). Due to the limited sample size, we are underpowered to deconvolute the effects of prior treatment history. Additionally, primary tumor genotyping information (n=4) or serial blood draws (n=11) were available in a small fraction of the total cohort. Finally, our benign group had a higher rate of *KRAS* codon 12 mutations than reported for similar molecular assays. We believe this is because we did not exclude patients with prior history of polyps or inflammation, who comprised 76% of benign samples tested. Though common in aging populations (26), these conditions are typically exclusion criteria in other ctDNA studies (8,31). However, we do not have tissue genotyping information from these patients to confirm this is the source of our “false positives”. Future studies, including matched tumor and ctDNA biopsies in an expanded patient cohort, are warranted to validate this assay for detection of localized PDA. Serial monitoring studies in patients with similar treatment histories will better inform the utility of this approach for PDA surveillance and early detection of relapse.

In summary, the data presented here suggest that MED-Amp could be a powerful screening tool for the detection of ctDNA at the earliest stages of PDA. An advantage of MED-Amp over other high-sensitivity strategies, such as allelic discrimination, is this method does not rely on selective blocking or degradation of wild-type alleles to enhance mutant template signal (32–34). Instead, by co-amplifying both wild-type and mutant-alleles in separate reaction volumes, both mutation abundance and frequency are preserved. Single molecule template emulsification also minimizes PCR bias associated with DNA fragment size and sequence motifs reported to suppress amplification (23). This strategy does not require laborious library construction or rigorous bio-statistical analyses to filter false-positives as with NGS-based methods. Robust performance at ultra-low DNA inputs means MED-Amp can also be adapted for a variety of challenging biological specimens. Expanding MED-Amp to include probes targeting a small panel of cancer-specific hotspots or incorporating additional biomarker data from routine patient follow up, could increase assay sensitivity with minimal increase in total cost and processing time. This strategy for rapid and low-cost genotyping could support existing clinical methods for biomarker-based risk stratification and therapy selection.

## Authors’ Contributions

**Conception and design:** E. D. Pratt, D. B. Zhen, A. D. Rhim

**Acquisition of data (provided animals, acquired and managed patients,provided facilities, etc.):** E. D. Pratt, D. B. Zhen, S. L. Manning, H. Cameron, K. Schradle, R.W. Cowan, D. M. Simeone, V. Sahai

**Analysis and interpretation of data (e.g., statistical analysis, biostatistics, computational analysis):** E. D. Pratt, A. D. Rhim

**Writing, review, and/or revision of the manuscript:** E. D. Pratt, D. B. Zhen, R. W. Cowan, A.D. Rhim

**Administrative, technical, or material support (i.e., reporting or organizing data, constructing databases:** V. Gunchick, H. Cameron, K. Schradle, V. Sahai, A. D. Rhim

