## Supplementary Information for "Ultra-low Input Circulating Tumor DNA Detection by MED-Amp in Early-Stage Pancreatic Cancer"

Table of Contents

| | |
|---|---|
| Table S-1 | Patient details for PDA patients |
| Table S-2 | Benign Patient Characteristics |
| Table S-3 | MED-Amp results for all patient sample |
| Table S-4 | Correlation between VAF fold change in time course plasma samples and PFS |
| Figure S-1 | Effect of tumor and sample storage characteristics on ctDNA detection |
| Figure S-2 | cfDNA input by stage and run-to-run sample variation |
| Figure S-3 | Survival analysis for patients who underwent a curative resection |

**Table S1: Patient details for PDA patients**

| ID | Disease | Stage | Location | Age | Gender | Tumor Size (cm) | Neoadjuvant | Adjuvant | Radiation | PFS (days) | PFS Status | OS (days) | OS Status |
|---|---|---|---|---|---|---|---|---|---|---|---|---|---|
| 157 | PDA | Metastatic | head | 59 | F | 3 | N | N | Y | | 0 | 1353 | 1 |
| 224 | PDA | Resectable | head | 76 | M | 2.3 | N | Y | Y | | 0 | 1562 | 0 |
| 278 | PDA | Metastatic | head /neck | 73 | F | 1.8 | N | N | Y | 87 | 1 | 1493 | 0 |
| 307 | PDA | Borderline | head | 71 | F | 4 | Y | N | Y | | 0 | 592 | 0 |
| 324 | PDA | Borderline | head | 50 | F | 1.4 | Y | N | Y | 102 | 1 | 426 | 0 |
| 342 | PDA | Resectable | head | 63 | M | No info | N | Y | Y | | 0 | 1418 | 0 |
| 386 | PDA | Locally Advanced | neck | 55 | F | 2.3 | N | Y | Y | 7 | 1 | 758 | 1 |
| 388 | PDA | Metastatic | head | 59 | M | 3.2 | Y | N | Y | 250 | 1 | 803 | 1 |
| 407 | PDA | Locally Advanced | body | 58 | F | 3.1 | N | N | N | 95+ | 1 | 435 | 0 |
| 427 | PDA | Metastatic | tail | 65 | M | 6.5 | N | N | N | | 0 | 436 | 0 |
| 483 | PDA | Borderline | neck | 81 | M | 3 | Y | N | N | No info | | 862 | 0 |
| 491 | PDA | Locally Advanced | neck | 63 | F | No info | Y | N | Y | 465 | 1 | 1242 | 0 |
| 510 | PDA | Metastatic | head | 46 | F | 2.8 | N | N | N | 66 | 1 | 1220 | 0 |
| 512 | PDA | Locally Advanced | body | 59 | F | No info | N | N | N | | 0 | 729 | 1 |
| 513 | PDA | Locally Advanced | head | 58 | F | 3.7 | N | N | N | 143 | 1 | 550 | 1 |
| 514 | PDA | Locally Advanced | body | 84 | F | 2.6 | Y | Y | N | 56 | 1 | 377 | 1 |
| 529 | PDA | Metastatic | head | 66 | F | 5.1 | N | N | Y | 273 | 1 | 735 | 1 |
| 530 | PDA | Borderline | head | 53 | M | 2.8 | Y | N | Y | | 0 | 1467 | 0 |
| 541 | PDA | Borderline | body / tail | 86 | F | 3.5 | Y | Y | Y | 652 | 1 | 1019 | 0 |
| 549 | PDA | Borderline | uncinate | 63 | M | 2.2 | Y | N | N | 320 | 1 | 641 | 1 |
| 557 | PDA | Borderline | neck / head | 65 | F | 2.1 | Y | Y | N | | 0 | 1090 | 1 |
| 558 | PDA | Resectable | head | 78 | M | 1.9 | N | N | N | | 0 | 619 | 1 |
| 560 | PDA | Borderline | head | 65 | F | 2.3 | Y | No info | N | No info | | 237 | 0 |
| 566 | PDA | Locally Advanced | head | 65 | M | 1.9 | N | Y | N | 424 | 1 | 734 | 1 |

| | | | | | | | | | | | | | |
|---|---|---|---|---|---|---|---|---|---|---|---|---|---|
| 570 | PDA | Locally Advanced | body | 73 | F | 4.3 | N | N | Y | 337 | 1 | 726 | 0 |
| 571 | PDA | Metastatic | body | 68 | F | No info | N | N | N | 266 | 1 | 501 | 1 |
| 576 | PDA/ Ampullary Cancer | Resectable | head | 58 | F | 2.5 | N | Y | Y | | 0 | 355 | 0 |
| 596 | PDA | Locally Advanced | neck / body | 65 | M | 6.7 | No info | No info | No info | No info | | 101 | 1 |
| 599 | PDA | Locally Advanced | head | 74 | F | No info | Y | N | Y | | 0 | 212 | 0 |
| 607 | PDA | Resectable | head | 61 | M | 2.6 | N | Y | N | 282 | 1 | 533 | 1 |
| 609 | PDA | Borderline | body | 57 | F | 4 | Y | N | Y | | 0 | 1159 | 0 |
| 611 | PDA | Metastatic | tail | 72 | F | 6.5 | N | N | N | | 0 | 429 | 0 |
| 628 | PDA | Metastatic | body | 76 | M | 2 | No info | No info | No info | No info | | 6 | 0 |
| 645 | PDA | Borderline | head | 53 | F | 2.5 | Y | N | Y | 787 | 1 | 1029 | 0 |
| 647 | PDA | Resectable | head | 73 | F | 2.6 | N | Y | Y | No info | | 398 | 1 |
| 662 | PDA | Borderline | tail | 57 | M | 4.7 | Y | Y | N | 573+ | 1 | 1070 | 0 |
| 665 | PDA | Metastatic | tail | 65 | F | 3.3 | N | N | N | | 0 | 308 | 1 |
| 670 | PDA | Locally Advanced | head | 74 | F | 2 | N | Y | N | 162 | 1 | 499 | 1 |
| 672 | PDA | Metastatic | body and tail | 47 | F | 4 | N | Y | N | | 0 | 258 | 1 |
| 673 | PDA | Borderline | head | 68 | M | 2.5 | Y | Y | N | 789 | 1 | 827 | 0 |
| 674 | PDA | Locally Advanced | head and body | 61 | F | 2.2 | N | Y | Y | | 0 | 203 | 1 |
| 677 | PDA | Metastatic | body | 63 | M | 4.4 | N | N | N | No info | | 151 | 1 |
| 682 | PDA | Borderline | body | 56 | F | 4.9 | N | N | Y | No info | | 270 | 0 |
| 684 | PDA | Metastatic | head | 65 | M | 3.6 | N | N | Y | 84 | 1 | 298 | 0 |
| 688 | PDA | Borderline | head/neck | 73 | M | 2.6 | Y | N | N | | 0 | 191 | 1 |
| 701 | PDA | Resectable | head | 62 | F | No info | N | Y | N | No info | | 48 | 0 |
| 703 | PDA | Resectable | tail | 68 | F | 2.3 | N | Y | No info | No info | | 556 | 0 |
| 734 | PDA | Locally Advanced | head | 81 | M | 2.8 | N | N | No info | No info | | 175 | 0 |
| 741 | PDA | Locally Advanced | head | 64 | F | 3.6 | No info | No info | No info | No info | | 94 | 1 |
| 742 | PDA | Resectable | body | 74 | F | 1.2 | N | N | N | No info | | 866 | 0 |

| | | | | | | | | | | | | | |
|---|---|---|---|---|---|---|---|---|---|---|---|---|---|
| 743 | PDA | Resectable | head | 72 | M | 2.9 | N | Y | N | | 0 | 907 | 0 |
| 745 | PDA | Metastatic | head | 61 | M | 4.1 | N | N | N | No info | | 69 | 0 |
| 747 | PDA | Metastatic | head | 59 | M | 3.1 | N | Y | N | 471 | 1 | 483 | 0 |
| 751 | PDA | Locally Advanced | head | 63 | M | 3 | No info | No info | No info | No info | | 26 | 0 |
| 753 | PDA | Metastatic | head | 75 | M | 4 | No info | No info | No info | No info | | 301 | 0 |
| 759 | PDA | Locally Advanced | body / neck | 62 | F | 2.6 | No info | No info | No info | No info | | 14 | 0 |
| 766 | PDA | Resectable | tail | 63 | F | 4 | N | Y | Y | 263 | 1 | 871 | 0 |
| 772 | PDA | Borderline | head / neck | 69 | F | 2.5 | Y | No info | Y | No info | | 301 | 0 |
| 773 | PDA | Borderline | head | 69 | M | 2.9 | Y | N | Y | | 0 | 862 | 0 |
| 774 | PDA | Locally Advanced | head | 80 | M | 4 | N | N | Y | 57 | 1 | 208 | 0 |
| 775 | PDA | Borderline | neck | 63 | M | 2.5 | Y | N | N | | 0 | 409 | 1 |
| 776 | PDA | Metastatic | head | 81 | M | 2 | N | Y | N | 224 | 1 | 459 | 1 |
| 780 | PDA | Resectable | head | 46 | M | 1.2 | N | N | N | | 0 | 840 | 0 |
| 784 | PDA | Metastatic | body/tail | 61 | F | 7.3 | N | N | N | 464 | 1 | 621 | 0 |
| 785 | PDA | Metastatic | body/tail | 59 | M | 5 | N | N | N | 222 | 1 | 548 | 1 |
| 800 | PDA | Locally Advanced | head | 81 | M | 2 | N | N | N | No info | | 668 | 0 |
| 801 | PDA | Resectable | head | 78 | M | 3 | N | N | N | No info | | 523 | 0 |
| 802 | PDA | Resectable | head | 63 | M | 2.2 | N | Y | N | No info | | 301 | 0 |
| 805 | PDA | Resectable | head | 75 | F | 1.6 | Y | N | Y | No info | | 204 | 0 |
| 814 | PDA | Resectable | Body | 74 | M | 2 | N | Y | N | No info | | 265 | 0 |
| 817 | PDA | Locally Advanced | Body/ neck | 79 | F | 6.1 | N | N | N | | 0 | 75 | 1 |
| 819 | PDA | Metastatic | body / tail | 62 | M | 7.5 | N | N | N | | 0 | 360 | 1 |

**OS:** median overall survival; **PDA:** Pancreatic Adenocarcinoma; **PFS:** median progression-free survival

**Table S2: Benign Patient Characteristics**

| Characteristics | Total n=25 | |
|---|---|---|
| **Age, median (range), yrs** | 54 | (51-58) |
| **Gender, no. (%)** | | |
| Female | 10 | (40.0) |
| Male | 15 | (60.0) |
| **Prior History, no. (%)** | | |
| Polyps | 11 | (44.0) |
| Inflammation | 3 | (12.0) |
| Polyps & Inflammation | 5 | (20.0) |
| Neither | 6 | (24.0) |

**Table S3: MED-Amp results for all patient samples**

| ID | Stage | Plasma Vol. (mL) | Variant Allelic Frequency (%) | No. MT Droplets | Estimated MT Copies | Detected by MED-Amp? |
|---|---|---|---|---|---|---|
| 819 | Metastatic | 2 | 13.9 | 1772 | 30 | Y |
| 628 | Metastatic | 2 | 12.96 | 24506 | 415 | Y |
| 665 | Metastatic | 2 | 6.67 | 46883 | 795 | Y |
| 611 | Metastatic | 2 | 4 | 7045 | 119 | Y |
| 775 | Borderline | 2 | 0.065 | 24.5 | 0 | N |
| 427 | Metastatic | 2 | 2.52 | 3849 | 65 | Y |
| 672 | Metastatic | 2 | 2.35 | 186 | 3 | Y |
| 157 | Metastatic | 2 | 2.16 | 3192 | 54 | Y |
| 814 | Resectable | 2 | 0.05 | 9 | 0 | N |
| 278 | Metastatic | 2 | 1.23 | 337 | 6 | Y |
| 734 | Locally Advanced | 2 | 0.1 | 33 | 1 | N |
| 759 | Locally Advanced | 2 | 0.08 | 17 | 0 | N |
| 766 | Resectable | 2 | 0.02 | 10 | 0 | N |
| 776 | Metastatic | 2 | 0.67 | 2047 | 35 | Y |
| 645 | Borderline | 1.5 | 0.06 | 33 | 1 | N |
| 780 | Resectable | 2 | 0.02 | 3 | 0 | N |
| 753 | Metastatic | 2 | 0.48 | 193 | 3 | Y |
| 745 | Metastatic | 2 | 0.36 | 205 | 3 | Y |
| 630 | Adenoma/Inflammation | 2 | 0.28 | 378 | 6 | Y |
| 571 | Metastatic | 2 | 0.2 | 152 | 3 | Y |
| 677 | Metastatic | 2 | 0.19 | 86 | 1 | Y |
| 560 | Borderline | 2 | 0.02 | 2 | 0 | N |
| 584 | Healthy | 2 | 0.17 | 59 | 1 | Y |
| 510 | Metastatic | 2 | 0.16 | 102 | 2 | Y |
| 785 | Metastatic | 2 | 0.16 | 7 | 0 | N |
| 670 | Locally Advanced | 2 | 0.05 | 31 | 1 | N |
| 805 | Resectable | 2 | 0.01 | 13 | 0 | N |
| 647 | Resectable | 2 | 0.005 | 4 | 0 | N |
| 570 | Locally Advanced | 2 | 0.01 | 6 | 0 | N |
| 558 | Resectable | 2 | 0 | 3 | 0 | N |
| 622 | Adenoma/Inflammation | 2 | 0.09 | 110 | 2 | Y |
| 566 | Locally Advanced | 2 | 0 | 1 | 0 | N |
| 751 | Locally Advanced | 2 | 0 | 4 | 0 | N |
| 614 | Adenoma/Inflammation | 2 | 0.07 | 110 | 2 | Y |
| 774 | Locally Advanced | 2 | 0 | 0 | 0 | N |
| 673 | Borderline | 2 | 0.02 | 8 | 0 | N |
| 741 | Locally Advanced | 2 | 0.88 | 1347 | 23 | Y |
| 604 | Adenoma/Inflammation | 2 | 0.06 | 51 | 1 | Y |
| 307 | Borderline | 2 | 0.01 | 26 | 0 | N |
| 743 | Resectable | 2 | 0 | 2 | 0 | N |

| | | | | | | |
|---|---|---|---|---|---|---|
| 639 | Adenoma/Inflammation | 2 | 0.05 | 92 | 2 | Y |
| 800 | Locally Advanced | 2 | 0.84 | 125 | 2 | Y |
| 802 | Resectable | 2 | 0 | 8 | 0 | N |
| 514 | Locally Advanced | 2 | 0.13 | 220 | 4 | Y |
| 674 | Locally Advanced | 2 | 0.09 | 207 | 4 | Y |
| 529 | Metastatic | 2 | 0.04 | 105 | 2 | Y |
| 817 | Locally Advanced | 2 | 0.07 | 82 | 1 | Y |
| 655 | Adenoma/Inflammation | 2 | 0.04 | 38 | 1 | Y |
| 684 | Metastatic | 2 | 0.04 | 3 | 0 | N |
| 801 | Resectable | 2 | 1.39 | 115 | 2 | Y |
| 773 | Borderline | 2 | 0.01 | 3 | 0 | N |
| 541 | Borderline | 2 | 0 | 8 | 0 | N |
| 609 | Borderline | 2 | 0 | 0 | 0 | N |
| 388 | Metastatic | 2 | 0.02 | 14 | 0 | N |
| 772 | Borderline | 2 | 0 | 4 | 0 | N |
| 662 | Borderline | 2 | 3.77 | 5701 | 97 | Y |
| 607 | Resectable | 2 | 0.71 | 931 | 16 | Y |
| 342 | Resectable | 2 | 0.49 | 463 | 8 | Y |
| 784 | Metastatic | 2 | 0.02 | 4 | 0 | N |
| 701 | Resectable | 2 | 0.11 | 116 | 2 | Y |
| 324 | Borderline | 2 | 0.54 | 50 | 1 | Y |
| 599 | Locally Advanced | 2 | 0.06 | 114 | 2 | Y |
| 682 | Borderline | 2 | 0.19 | 116 | 2 | Y |
| 386 | Locally Advanced | 2 | 0.04 | 40 | 1 | Y |
| 512 | Locally Advanced | 2 | 0.04 | 39 | 1 | Y |
| 530 | Borderline | 2 | 0.03 | 45 | 1 | Y |
| 703 | Resectable | 2 | 0.1 | 138 | 2 | Y |
| 224 | Resectable | 2 | 0.09 | 138 | 2 | Y |
| 557 | Borderline | 2 | 0.03 | 62 | 1 | Y |
| 742 | Resectable | 2 | 0.06 | 43 | 1 | Y |
| 596 | Locally Advanced | 2 | 0.04 | 53 | 1 | Y |
| 583 | Adenoma/Inflammation | 2 | 0 | 2 | 0 | N |
| 589 | Healthy | 2 | 0 | 3 | 0 | N |
| 593 | Adenoma/Inflammation | 2 | 0 | 2 | 0 | N |
| 602 | Adenoma/Inflammation | 2 | 0 | 3 | 0 | N |
| 603 | Adenoma/Inflammation | 2 | 0 | 2 | 0 | N |
| 605 | Adenoma/Inflammation | 2 | 0 | 0 | 0 | N |
| 606 | Adenoma/Inflammation | 2 | 0 | 1 | 0 | N |
| 688 | Borderline | 1.5 | 0.03 | 186 | 3 | Y |
| 624 | Adenoma/Inflammation | 1 | 0 | 2 | 0 | N |
| 625 | Adenoma/Inflammation | 2 | 0 | 5 | 0 | N |
| 633 | Healthy | 2 | 0 | 6 | 0 | N |
| 634 | Healthy | 2 | 0 | 2 | 0 | N |
| 635 | Adenoma/Inflammation | 2 | 0 | 21 | 0 | N |
| 636 | Adenoma/Inflammation | 2 | 0 | 0 | 0 | N |

| | | | | | | |
|---|---|---|---|---|---|---|
| 637 | Healthy | 2 | 0 | 33 | 1 | N |
| 638 | Adenoma/Inflammation | 2 | 0 | 16 | 0 | N |
| 649 | Adenoma/Inflammation | 2 | 0 | 3 | 0 | N |
| 654 | Healthy | 2 | 0 | 1 | 0 | N |
| 659 | Adenoma/Inflammation | 2 | 0 | 1 | 0 | N |
| 696 | Resectable | 2 | 0.04 | 84 | 1 | **Y** |
| 747 | Metastatic | 2 | 0 | 5 | 0 | N |
| 407 | Locally Advanced | 2 | 0.01 | 91 | 2 | **Y** |
| 483 | Borderline | 2 | 0.01 | 53 | 1 | **Y** |
| 513 | Locally Advanced | 2 | 0.01 | 93 | 2 | **Y** |
| 576 | Resectable | 2 | 0.015 | 63 | 1 | **Y** |

**MT:** Mutant *KRAS*+ (G12 C/D/R/V)

**Table S4: Correlation between VAF fold change in time course plasma samples and progression-free survival**

| ID | Stage | Draw 1 | | | Draw 2 | | | Days Between Draws | VAF Fold Change | VAF Directional Change | PFS Status |
|---|---|---|---|---|---|---|---|---|---|---|---|
| | | Allelic Frequency (%) | No. MT Droplets Detected | Detected by MED-Amp? | Allelic Frequency (%) | No. MT Droplets Detected | Detected by MED-Amp? | | | | |
| **278** | Metastatic | 0.47 | 337 | Y | 1.23 | 716 | Y | 101 | 1.62 | Up | 1 |
| **491** | Locally Advanced | 0.03 | 145 | Y | 0.1 | 248 | Y | 129 | 2.33 | Up | 1 |
| **513** | Locally Advanced | 0.01 | 93 | Y | 0 | 5 | N | 91 | -1 | Down | 1 |
| **570** | Locally Advanced | 0.01 | 6 | N | 0.02 | 22 | N | 113 | 1 | Up | 1 |
| **571** | Metastatic | 0.2 | 152 | Y | 1.95 | 315 | Y | 267 | 8.75 | Up | 1 |
| **684** | Metastatic | 0.04 | 3 | N | 0.24 | 3 | N | 98 | 5 | Up | 1 |
| **157** | Metastatic | 2.16 | 3192 | Y | 2.4 | 77 | Y | 91 | 0.11 | No Change | 0 |
| **224** | Resectable | 0.09 | 138 | Y | 0 | 8 | N | 126 | -1 | Down | 0 |
| **530** | Borderline | 0.03 | 45 | Y | 0.17 | 96 | Y | 105 | 4.67 | Up | 0 |
| **557** | Borderline | 0.03 | 62 | Y | 0 | 0 | N | 140 | -1 | Down | 0 |
| **674** | Locally Advanced | 0.09 | 207 | Y | 0 | 2 | N | 112 | -1 | Down | 0 |
| **560** | Borderline | 0.02 | 2 | N | 0 | 7 | N | 147 | -1 | Down | -- |
| **682** | Borderline | 0.19 | 116 | Y | 0.01 | 47 | Y | 107 | -0.95 | Down | -- |
| **703** | Resectable | 0.1 | 138 | Y | 0.1 | 106 | Y | 63 | 0 | No Change | -- |

**MT:** Mutant *KRAS*+ (G12 C/D/R/V); **PFS:** Progression-Free survival; **VAF:** Variant Allelic Frequency

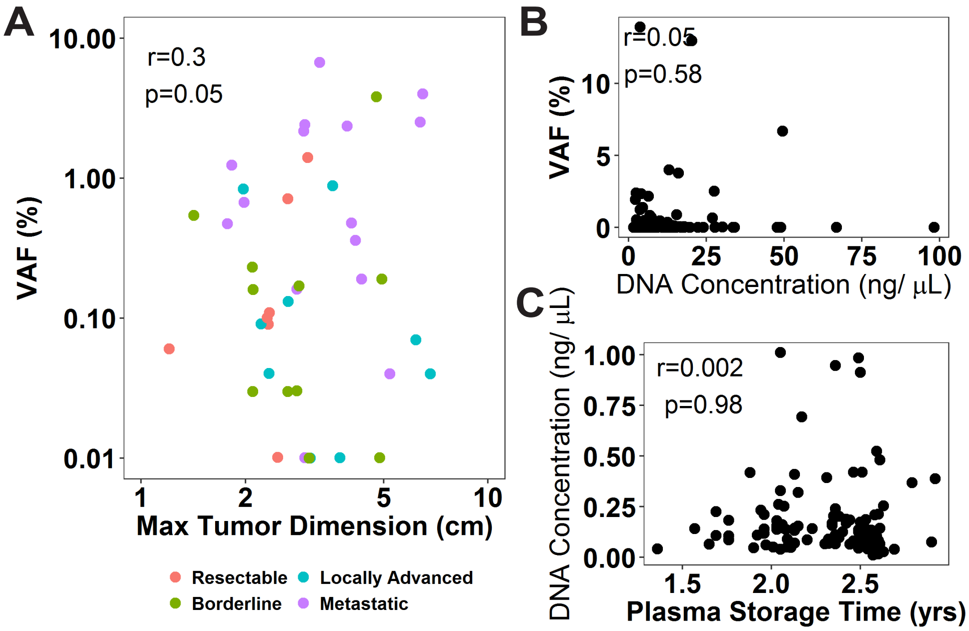

**Figure S-1.** Effect of tumor and sample storage characteristics on ctDNA detection. A) Pearson correlation between max tumor dimension (cm) and measured ctDNA allelic frequency. Colors indicate PDA stage at diagnosis. Correlation between B) concentration of cfDNA isolated from plasma samples and measured ctDNA VAF, and C) plasma storage time and amount of isolated cfDNA.

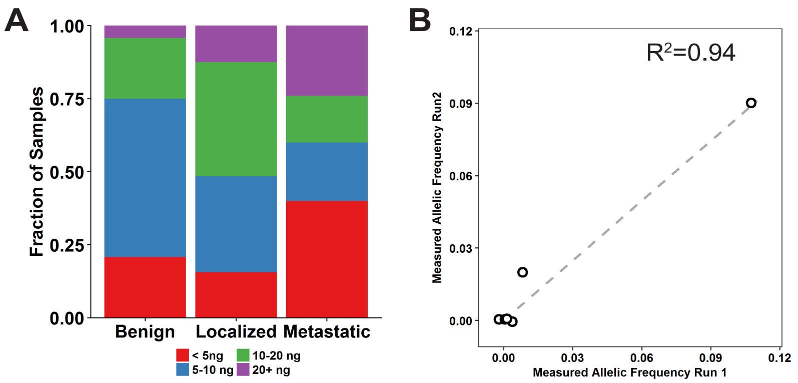

**Figure S-2.** A) Distribution of cfDNA inputs for MED-Amp analysis. B) Linear regression of concordance between two independent MED-Amp runs of an identical plasma sample.

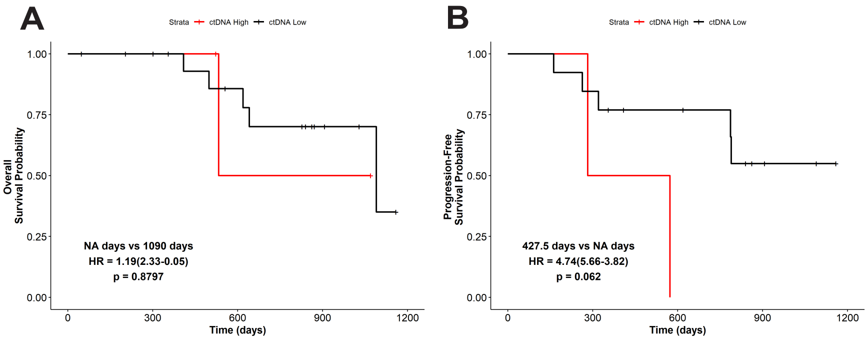

**Figure S-3.** Survival analysis of patients who underwent a curative resection shows pre-surgery ctDNA did not predict OS (A) or PFS (B).
